# In vivo tracking of CAR-T cells in tumors via nanobubble-based contrast enhanced ultrasound

**DOI:** 10.1101/2025.05.06.651695

**Authors:** Dorian Durig, Jude Franklin, Reshani Perera, Zachary Jackson, Hosahalli Vasanna, Michael C. Kolios, David Wald, Agata A. Exner

## Abstract

CAR-T cell therapy has led to remarkable advances in the outcomes of patients with acute lymphoblastic leukemia (ALL), B cell lymphomas, and multiple myeloma. Given these successes in hematologic malignancies, extensive efforts are now focused on developing CAR-T cell therapies for the treatment of solid tumors. The treatment of solid tumors poses significant hurdles with cell trafficking that is necessary to achieve efficacy and minimize off-tumor side effects. The development of simple, safe and inexpensive modalities to track CAR-T cell distribution in humans could provide critical insights to facilitate the development of improved CAR-T products for solid tumors. Here we demonstrate a strategy to monitor CAR-T cells *in vivo* using ultrasound imaging and nanobubble (NB) labeled cells. NBs are ultrasound contrast agents composed of a lipid shell and a C_4_F_10_ gas core that can be efficiently internalized into cells. This approach enables us to image the CAR-T cells using nonlinear contrast-enhanced ultrasound (CEUS). Utilizing this method, we found that CAR-T cells can be visualized after injection into both tumor bearing and non-tumor bearing mice. In summary, our ultrasound-based tracking approach can effectively monitor the trafficking of CAR-T cells *in vivo*, offering a valuable new strategy that can further enable the development of new CAR-T products and strategies to modulate cell trafficking.

## Introduction

Chimeric antigen receptor T cell (CAR-T) therapy is an advanced form of adoptive cell therapy (ACT) that has significantly impacted the field of immunotherapy. CAR-T cell therapy uses patient-specific T cells to create several FDA-approved products to treat B-cell malignancies and multiple myeloma.^1,2^ The manufacturing process involves the genetic engineering of the patient’s T cells to express a chimeric antigen receptor directed towards a target antigen expressed on tumor cells such as CD19, which is expressed specifically on malignant and normal B cells. The genetically modified T cells are intravenously infused into the patient where it is hoped that they will traffic to the tumor, expand and kill the tumor cells.

Despite notable achievements in hematological malignancies, the success of CAR-T therapy has not been reproduced in treating solid tumors.^3^ The limited success is hypothesized to be attributed to multiple factors, including an immunosuppressive tumor microenvironment, on-target off-tumor toxicity, T cell exhaustion, tumor heterogeneity, poor trafficking to the tumors, and lack of CAR-T cell persistence.^4^ To understand the factors contributing to the limited success of CAR-T cells in treating tumors and better finetune dosage parameters while mitigating side effects, such as neurotoxicity and cytokine release syndrome, the biodistribution and trafficking of the cells to the target after infusion need to be studied.^5^

Currently, clinically available methods are inadequate for assessing the real-time trafficking patterns of CAR-T cells in organs and tumors, which is essential for evaluating treatment efficacy and adjusting strategies accordingly. The standard practice for monitoring cellular kinetics and patient responses to immunotherapy involves intermittent blood draws, which provides valuable information such as white blood cell counts and the presence of tumor DNA.^6^ However, these tests fail to reveal the biodistribution and movement of CAR-T cells within the body. Real-time tracking of immune treatments would allow for a more comprehensive understanding of how CAR-T cells engage with the tumor microenvironment, providing timely insights that can improve therapeutic outcomes.

The benefits of tracking CAR-T cells via imaging have recently been recognized, leading to the adoption of various techniques that utilize modalities such as MRI, PET, and optical imaging, each providing unique information that surpasses the periodic blood draws used in the clinic.^7-12^ These approaches, however, also have disadvantages and limitations: MRI suffers from low temporal resolution and high costs, PET is associated with radiation exposure and limited spatial resolution, while optical imaging has shallow penetration depth and is restricted to superficial tissues.^13,14^ Although each technique offers valuable information, no single method has been established as the definitive standard for monitoring the trafficking and distribution of cells. To provide an alternative strategy for tracking CAR-T cells in vivo, we investigate using contrast-enhanced ultrasound (US).

US imaging is a non-ionizing modality that is widely used and accessible at a low cost. It offers deep penetration and high spatial and temporal resolution, making it extensively used for ultrasound-guided procedures and examinations, including guiding cell transplant injections.^15-18^ However, even in these applications, the transplanted cells can be difficult to differentiate from surrounding soft tissue, mirroring the challenges faced in identifying immune cells following immunotherapy treatment. As a result, US contrast agents (UCAs) have become a promising tool for enhancing cell visibility by labeling cells, thereby increasing their contrast relative to surrounding tissue. ^18-20^

UCAs are made up of lipids, proteins, or polymers that form a shell around a gas core, creating a bubble.^21^ Microbubbles (MBs), which are commercially available UCAs, range in size from approximately 1 to 8 µm and are primarily used to enhance vascular imaging due to their confinement within the vasculature.^21,22^ However, nanosized contrast agents, such as nanobubbles (NBs), have emerged as an alternative to MBs due to their smaller size of approximately 100-600 nm in diameter.^15^ This smaller size allows NBs to move beyond the vasculature, making them a more promising UCA for labeling cells expected to extravasate and infiltrate tissue. Additionally, the size of NBs enables their internalization into cells, allowing them to serve as effective labels for tracking. Several studies have demonstrated the effectiveness of nanosized contrast agents in labeling and tracking various cell types, including stem cells and NK cells.^18,20,23-25^ Building on this foundation, we aim to track CAR-T cells using US imaging enhanced by NBs, which could enable timely and cost-effective monitoring of these therapeutic cells.

Prior studies have demonstrated the ability of NBs to be trafficked inside cells via phagocytosis or receptor-mediated endocytosis.^25,26^ We hypothesize that CAR-T cells may also utilize endocytosis to uptake NBs during co-incubation, enabling non-invasive, real-time monitoring of infused cells. To evaluate this technique, we labeled CD19-targeted CAR-T cells because of their widespread clinical use for Lymphoma and the potential of this monitoring method to enhance an approved immunotherapy (Fig. 1). We then assessed the trafficking of labeled CD19 CAR-T cells using CEUS in both a healthy murine model and a Burkitt’s lymphoma (RAJI cells) murine model. In healthy tissue, CAR-T cells exhibited rapid influx, post-injection, while in the tumor model, accumulation was notably slower. Our findings show that CD19 CAR-T cells rapidly traffic to and remain localized in the tumor for up to 75 minutes post-infusion. While therapeutic effects are typically observed at later time points, this early localization, observable due to our NB labeling, highlights the cells’ efficient homing to the tumor site immediately following infusion. These results demonstrate the feasibility of using this imaging technique for non-invasive, real-time tracking of CAR-T cells, underscoring its potential for clinical application in CAR-T cell therapy monitoring.

**Fig. 1.**
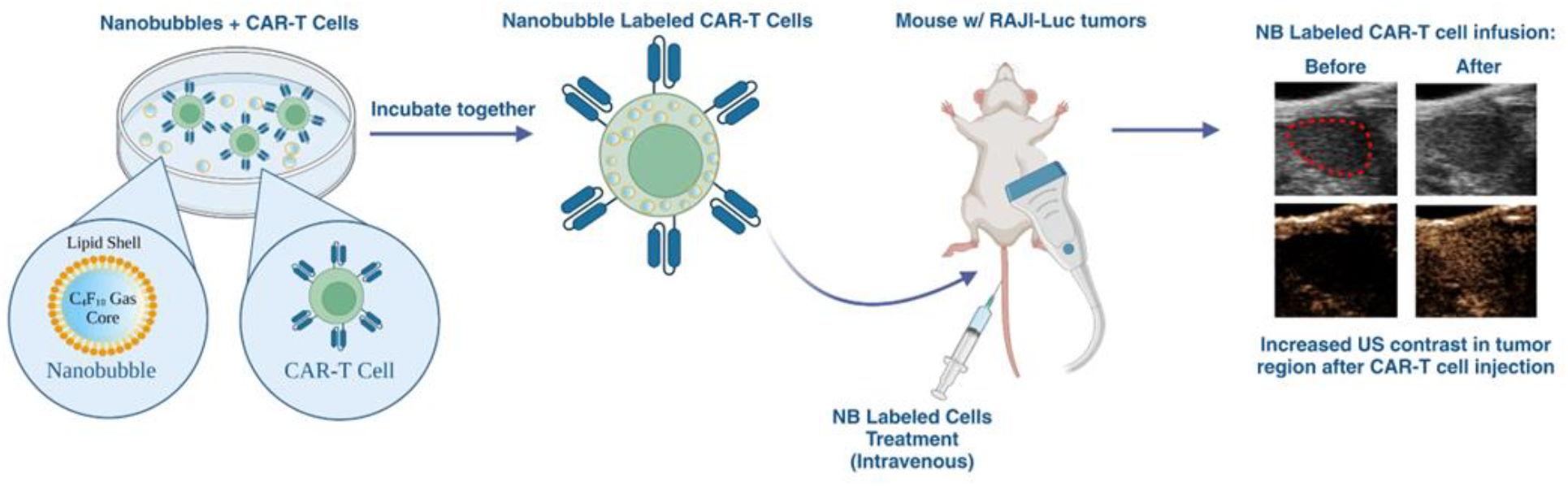
Schematic illustration of the NB labeling process of CAR-T cells and their injection into tumor bearing mice. Images are not to scale.

## Materials and Methods

### Preparation of CAR-T cells

Peripheral blood mononuclear cells (PBMCs) were isolated from healthy donor whole blood samples mononuclear peripheral blood samples were processed within 24 hours of receipt. The study was approved by the University Hospital Cleveland Medical Center institutional review board and all donors gave written informed consent. T cells were stimulated using Miltenyi TransAct reagent (Miltenyi Biotec; Bergisch Gladbach, Germany). In addition, T cells were supplemented with cytokines (IL2 every other day to a final concentration of 30IU/ µL). The activated T cells were cultured in complete RPMI medium supplemented with 10% fetal bovine serum (FBS; Cytiva; Marlborough, MA) and 1% Penicillin-streptomycin antibiotics (Pen Strep; Cytiva; Marlborough, MA).27 During labeling, however, the cells were cultured in 3% human serum albumin. Following activation, T cells were cultured with periodic media changes, maintaining a concentration of 1 million cells/mL.

For genetic modification, CD19 CAR Lentiviral particles were produced by transfecting HEK293T cells using Lipofectamine™ 3000 (Thermo Fisher Scientific; Waltham, MA) with a second generation 41BB/CD3zeta containing CAR plasmid and helper plasmids, encoding as VSV-G, GAG/POL, and REV (Addgene; Watertown, MA). The T cells were transduced with the corresponding CD19 lentiviral vector, 1 day after activation, and expanded for a maximum of 10 days before cryo-freezing the cells in liquid nitrogen.^28^

At day 5 and onwards, the expanded CAR-T cells were assessed for viability, CAR-expression, and cytotoxicity. The viability was assessed by trypan blue counts, and the CAR expression was assessed by flow cytometry using anti-FMC63 FITC (Acrobiosystems; Newark, DE) on the Attune Flow Cytometer (Thermo Fisher Scientific; Waltham, MA). In addition, the functionality of the CAR-T cells was validated for cytotoxicity via in vivo survival study.

### Preparation of nanobubbles

NBs were prepared according to a previously published protocol (de Leon et al).^29^ However, we used C_4_F_10_ as an alternative gas, which has shown to improve stability compared to C_3_F_8_.^30,31^ Briefly, the lipids DPPA (1,2-dipalmitoyl-sn-glycero-3-phosphate; Avanti Polar Lipids Inc.; Pelham, AL), DPPE (1,2-dipalmitoyl-sn-glycero-3-phospho-ethanolamine; Avanti Polar Lipids Inc.; Pelham, AL), DSPE-mPEG-2k (1,2-distearoyl-snglycero-3-phosphoethanolamine-N-[methoxy(polyethylene glycol)-2000]; Laysan Lipids; Arab, AL), and DBPC (1,2-dibehenoyl-snglycero-3-phosphocholine; Avanti Polar Lipids Inc.; Pelham, AL) were dissolved in propylene glycol (PG; Sigma Aldrich; Saint Louis, MO) at 80°C in a hot water bath. Simultaneously, a solution of phosphate-buffered saline (PBS; Gibco Life Technologies; Grand Island, NY) and glycerol (Sigma Aldrich; Saint Louis, MO) was heated to 80°C. Once the lipids were fully dissolved, the PBS-glycerol solution was added to the lipid mixture. The final solution was added to a 3 mL headspace vial and sealed. The gas was then exchanged with C_4_F_10_ (Perfluorobutane; FluoroMed; Round Rock, TX) and the vials were activated using mechanical agitation (Vialmix). NBs were isolated by centrifugation for 5 minutes at 50 g and extracted prior to use. Rhodamine-NBs were formulated by incorporating DSPE-PEG-Rhodamine (1,2-dioleoyl-sn-glycero-3-phosphoethanolamine-N-(lissamine rhodamine B sulfonyl); Avanti Polar Lipids Inc.; Pelham, AL) into the lipid mixture prior to lipid dissolution. The NBs were characterized using dynamic light scattering (DLS; Litesizer DLS 500, Anton Paar) demonstrating plain NBs with no fluorescent label have an intensity-weighted diameter of 293.52 ± 50 nm, while rhodamine-labeled NBs have an intensity-weighted diameter of 258.52 ± 42 nm (supplementary data). Based on previously characterized NB formulations using resonant mass measurement (RMM; Archimedes®, Malvern Panalytical), the NBs are assumed to have a concentration of 3.5 × 10^11^ NBs/mL.^24,29,32^

### Labeling CAR-T cells with nanobubbles

CD19 CAR-T cells were thawed in complete RPMI media containing 10% FBS and 1% Pen Strep. After thawing, the cells were left in the complete media for 30 minutes before being centrifuged for 5 minutes at 500 g and collected. The cells were then diluted to 5×10^6^ cells/mL in RPMI media with 3% human serum albumin (HSA; Innovative Research; Novi, MI). NBs were added to the cell solution to achieve concentrations of 10k, 15k, or 20k NBs/cell. The NB-cell solution was incubated at 37°C with 3% CO_2_ for 1 hour, with gentle shaking every 15 minutes. After the incubation period, the cells were centrifuged at 230 g for 10 minutes and washed in incomplete media.

### In vitro ultrasound imaging setup

The in vitro US imaging was conducted in a three well agarose phantom shown in Figure 2a. The phantom was prepared by dissolving low-temperature gelling agarose (Sigma Aldrich; Saint Louis, MO) at a 2.4% w/v concentration in deionized (DI) water. The agarose solution was then poured into a mold with the desired specifications (Fig. 2a). During imaging, the MS250 US transducer (Vevo® 2100, FUJIFILM VisualSonics) was positioned perpendicular to the wells against the phantom and coupled with US gel. Approximately 50 µL of the cell suspension was placed in the well and imaged using nonlinear contrast (NLC) mode with the following parameters: 4% power, 18 MHz, 1 fps, 35 dB contrast gain, 18 dB 2D gain. The nonlinear contrast mode on the Vevo 2100 uses a form of amplitude modulation to generate the images.^33^

**Fig. 2.**
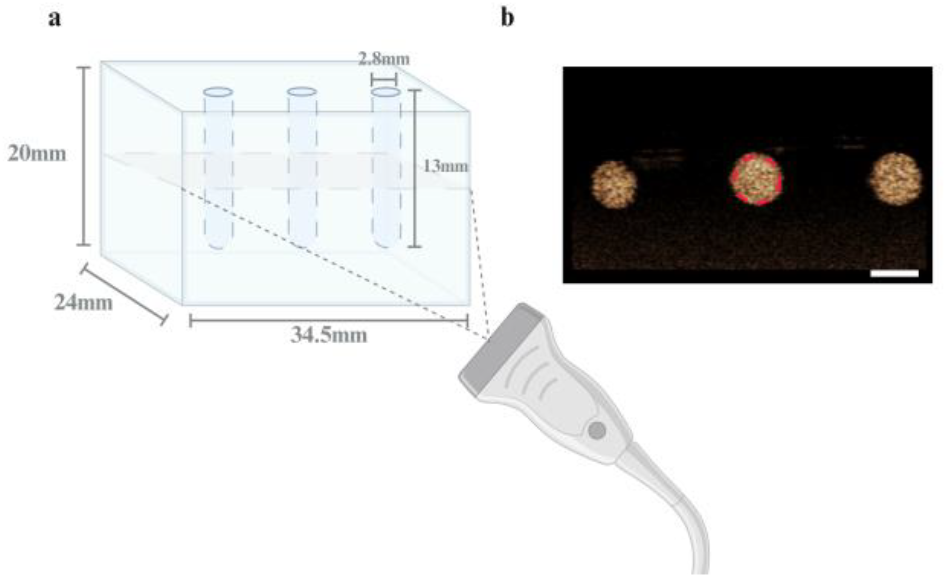
(a) In vitro imaging setup using the 3-well agarose phantom with the transducer placed horizontally at the side of the phantom. (b) CEUS cross-sectional image produced by the in vitro imaging setup using the phantom with all 3 wells filled with NB contrast agent. The red circle over the center well indicates the ROI used to analyze the average signal intensity (scale bar = 2.8mm).

### In vitro ultrasound imaging

NB-labeled CAR-T cells were evaluated in vitro utilizing two distinct methods within the US imaging setup: (a) Imaging the labeled cells at discrete time-points over 24 hours, and (b) Minimum detectable concentration. In the discrete time-point study, cells were labeled with 10k, 15k, and 20k NBs/cell. After labeling, the cells were diluted to 1×10^6^ cells/mL in complete RPMI media and transferred to a 96-well plate with 200 µL per well. The samples were imaged immediately after labeling, and then at 0.5, 1, 2, 3, 6, and 24 hours post-labeling. Between imaging sessions, the cells were kept in an incubator at 37°C with 5% CO_2_. To evaluate the minimum detectable concentration the cells were labeled with 20k NBs/cell. After labeling, the cells are diluted to 1×10^6^, 10^5^, 10^4^, and 10^3^ cells/mL in PBS and evaluated in the phantom. For each study the control samples containing only CAR-T cells underwent the same labeling conditions and imaging parameters, but without the addition of NBs.

### In vivo ultrasound imaging

Male and female NCG mice (strain code: 572; Charles River) were handled in according to a protocol approved by the Institutional Animal Care and Use Committee (IACUC) at Case Western Reserve University. The RAJI cells (ATCC) were injected intravenously (2×10^6^ cells in 150 µL of PBS) to establish disease of the course of three weeks. Tumor growth was monitored using bioluminescence imaging (PerkinElmer), US imaging was performed around the left kidney after confirmation of tumor growth in this region by bioluminescence imaging. Approximately 0.7-1×10^7^ NB labeled CAR-T cells were injected intravenously, and mice were imaged during and after labeled cell infusion at the following time points: 0, 5, 15, 30, 45, 60, and 75 minutes. Each time point had n=4, except for the 5-minute time point, where n=3. Liver imaging studies were conducted separately in non-tumor bearing NCG mice. The liver was imaged at the same time points as the tumor-bearing mice (n=3).

US imaging was conducted using the Vevo 2100 ultrasound system with the MS250 transducer in NLC mode. The same parameters were used as the in vitro studies: 4% power, 18 MHz, 1 fps, 35 dB contrast gain, and 18 dB 2D gain. All animals underwent hair removal in the region of interest (ROI) and were anesthetized via inhalation of 2% isoflurane mixed with 1.5 L/min of air and were placed on a heating platform to maintain a stable temperature throughout the imaging timeframe.

### Image analysis

NLC images were quantified using FUJIFILM VisualSonics Vevo LAB software. For in vitro data analysis, an ROI was drawn, as shown in Fig. 2b, with an additional ROI selected outside the well of the phantom to quantify the background signal. The background was subtracted from the average signal intensity within the well to provide comparability between samples. For the analysis of in vivo studies, ROIs were manually drawn over each area of interest (liver, kidney, and tumor) to exclude surrounding tissues. The baseline NLC signal in each tissue was acquired and quantified prior to the infusion of labeled cells. Following this background acquisition, the nonlinear signal intensity was measured for the infusion of labeled cells, with the maximum value used for the 0 min discrete time point. The mean nonlinear signal was then calculated for frames captured at 5, 15, 30, 45, 60, and 75 minutes post-infusion. The baseline signal was used to compute the signal-to-baseline ratio over the 75-minute period and to adjust the NLC signal over time by subtracting the background.

### Confirmation of CAR-T cell labeling

Labeling efficiency of CAR-T cells with Rhodamine-conjugated NBs was assessed through flow cytometry and microscopy for flow cytometry analysis, following the labeling procedure described in the methods. A minimum of 200,000 events per sample were collected to ensure robust data. Flow cytometry data were analyzed using Attune software and formalized with FlowJo software (BD Biosciences, NJ). Rhodamine-conjugated NB-labeled CD19 CAR-T cells were compared to unlabeled CD19 CAR-T cells, which served as controls to verify the labeling process. Uptake of Rhodamine-conjugated NBs by CAR-T cells was quantified through the positive shift in Rhodamine fluorescence, with distinct separation from the control group, confirming successful and efficient labeling.

CAR-T cell labeling was also confirmed by confocal microscopy. Here, CAR-T cells were labeled with rhodamine-labeled NBs and then diluted to 1×10^6^ – 2×10^6^ cells/mL in PBS. The cells were fixed onto a glass slide using 4% paraformaldehyde (PFA; Fisher Chemicals, Pittsburgh, PA) for 30 minutes, washed with PBS three times, stained using DAPI mounting media (Fisher Chemical; Pittsburgh, PA) and sealed with a glass coverslip. Slides were kept at 4°C until imaging. Imaging was performed on a Leica SP8 confocal microscope.

### In vitro assessment of viability and differentiation via flow cytometry

To assess CAR-T cell proliferation, both NB-labeled CAR-T cells and unlabeled control T cells were cultured and counted every two days following culture initiation. Cell counts were performed using the Countess (Invitrogen; Waltham, MA) with trypan blue exclusion to distinguish viable cells from non-viable ones.

To evaluate differentiation and viability following labeling, CAR-T cells were stained with the following reagents: CD3 (APC), CD4 (APCeflour780), CD8 (BV510), CD45RA (BV605), CCR7 (PE), CAR antigen (FITC), and CD27 (AF700) antibodies, as well as the viability dye 7-AAD), (BD BioSciences; Franklin Lakes, NJ). Flow cytometry data were analyzed using FlowJo software (BD Biosciences; Franklin Lakes, NJ). The memory panel, assessing differentiation markers, was performed 4 days after labeling to monitor any potential effects over time. Additionally, the study was verified across a range of NB concentrations (1k, 5k, 10k, and 20k NBs/cell) to determine if the labeling process had any concentration-dependent impact on CAR-T cell differentiation. For this study, CAR-T cells were labeled using C_3_F_8_ NBs. The labeling protocol consisted of a 1-hour incubation on a shaker set to 225 rpm (linear rocking). The cells were incubated in complete RPMI medium, with a seeding density of 5 × 10^6^ cells/mL, with the incubator maintained at 5% CO_2_.

### In vivo efficacy and biodistribution

An efficacy study was conducted to verify the in vivo functionality of the NB-labeled CAR-T cells compared to the unlabeled CAR-T cell control group. Male and female mice received intravenous injections via the tail vein with 1 million luciferase-expressing RAJI tumor cells. In the inoculated mice used for the US imaging, CAR-T cells are injected 1 week following tumor inoculation. Approximately 1×10^7^ CAR-T cells, labeled with 20k NBs/cell following the same protocol used in the differentiation study, were administered intravenously via the tail vein. Progression of the tumor burden was monitored weekly by weight measurements and bioluminescence imaging using the IVIS Spectrum Imager (PerkinElmer; Waltham, MA). Mice were injected with 10 mg/kg of D-Luciferin (Invitrogen; Waltham, MA) subcutaneously. After 10 minutes, the images were taken and then analyzed with Living Image (PerkinElmer). Background values were taken prior to luciferase imaging to remove noise from the bioluminescent regions of interest. Tumor burden was quantified in terms of luciferase intensity (radiance).

### Statistical analysis

NLC data was plotted and analyzed using Excel and GraphPad Prism, with each experiment conducted three times unless otherwise specified. Unpaired two-tailed t-tests were utilized to compare the two groups. Data is presented as mean ± SEM (standard error of the mean), unless stated otherwise, and a p-value of less than 0.05 was deemed statistically significant. Outliers were assessed and removed using Grubbs’ test with a significance threshold of p = 0.05.

## Results and Discussion

### In vitro evaluation of labeled CAR-T cells

The labeling efficiency of CAR-T cells with NBs was first evaluated in vitro using an agarose phantom (Fig.3a-b). Unlabeled control CAR-T cells showed negligible signal, with contrast values of 10^6^ cells/mL (19.05 ± 6.17 a.u.) and 10^3^ cells/mL (33.64 ± 2.40 a.u.). In comparison, NB-labeled CAR-T cells demonstrated significantly higher signals at 10^6^ cells/mL (64,032.52 ± 20,859.39 a.u.) and 10^3^ cells/mL (105.09 ± 60.02 a.u.), showing a 3,362-fold and 3-fold increase, respectively, over unlabeled cells.

Based on these results, the detection threshold for the US acquisition parameters used in these studies is 10^3^ cells/mL (lower concentrations were not evaluated). This demonstrates a high detection sensitivity compared to other techniques, which report thresholds between 10^4^ and 10^5^ cells/mL.^9,12^

To evaluate the stability of the signal within labeled cells and the effects of NB concentration on the labeling process, CAR-T cells were incubated at 37°C with 10k, 15k, and 20k NBs/cell and imaged at 0.5, 1, 2, 3, 6, and 24 hours (Fig.3c). All samples exhibited high contrast immediately after labeling. The signal gradually decreased to baseline over 24 hours (Fig. 3d). Cells labeled with 20k NBs/cell showed the highest average nonlinear signal at all time points. At 6 hours, the signal in these cells remained 5.5 times that of the unlabeled cells. The average signal for the 10k and 15k NB/cell samples decayed faster and returned to baseline at 6 hours. From this study we opted to use the 20k NBs/cell concentration for labeling CAR-T cells.

**Fig. 3.**
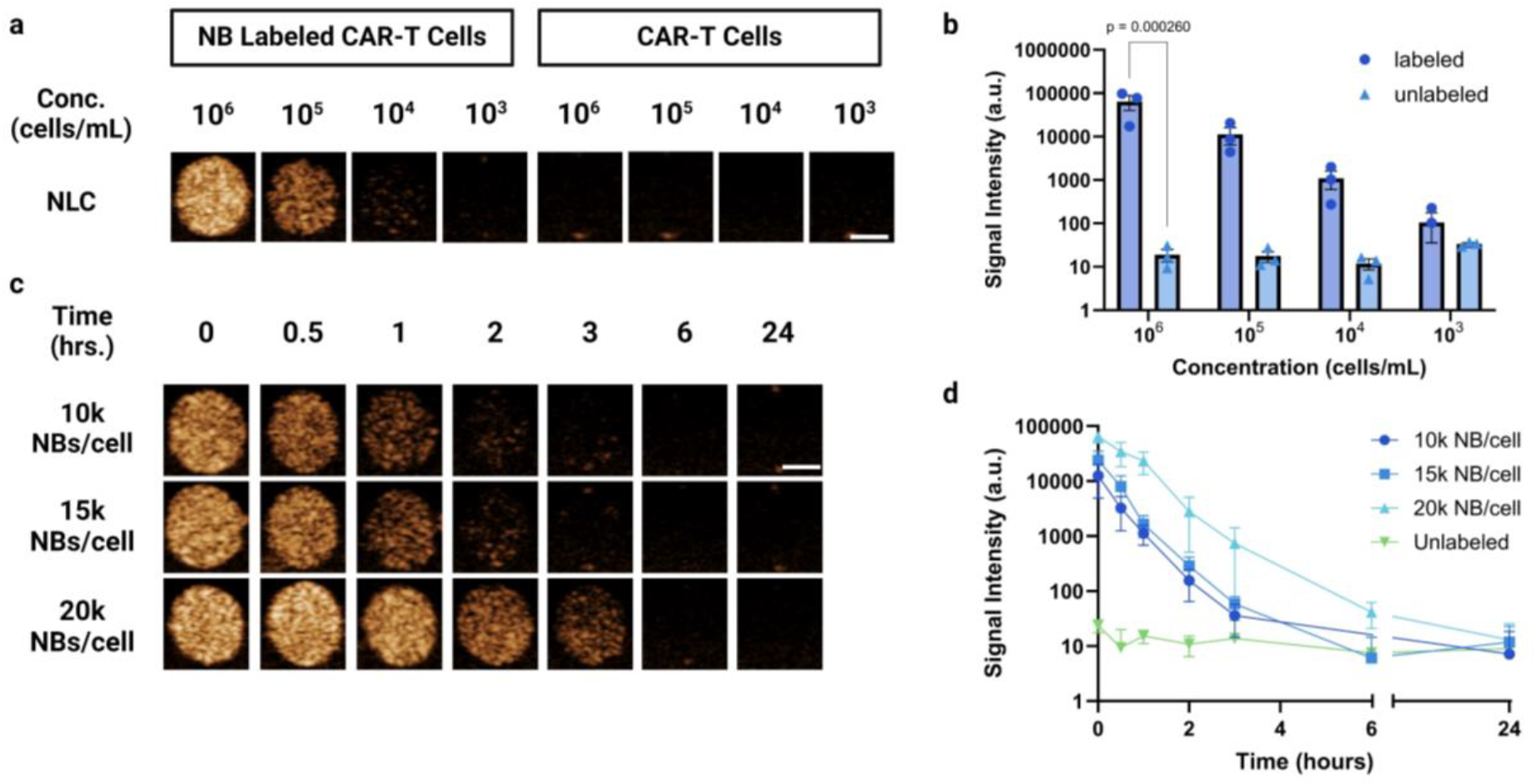
(a) CEUS images of CAR-T cells labeled with 20k NBs/ cells and unlabeled CAR-T cells at concentrations of 1×10^3^-1×10^6^ cells/ mL (scale bar = 1.4mm) (b) Average contrast in well at different concentrations of labeled (n=3) and unlabeled cells (n=3) (c) CAR-T cells labeled with various concentrations of NB per cell during the co-incubation process. (scale bar = 1.4mm) (d) Average contrast overtime of CAR-T cells labeled with 10k,15k, and 20k NBs/cell (n=3) and unlabeled cells (n=4; n=3 for 6hr).

### In vivo detection of labeled CAR-T cells in non-tumor bearing mice

Using the optimized concentration of 20k NBs/cell for labeling CAR-T cells, the CAR-T cells were evaluated in vivo through intravenous administration to immunodeficient mice (Fig. 4a). The liver was selected for initial evaluation due to its relatively low baseline signal with US. After infusion, the liver signal in the nonlinear mode increased by an average of 13-fold over baseline, followed by a gradual decrease over 75 minutes (Fig. 4b). This study serves as a proof-of-concept to demonstrate the feasibility of tracking CAR-T cells in vivo using US. It is likely that the signal detected during this time course represents CAR-T cells transiently passing through the liver as current research suggests that CAR-T cells begin to accumulate to higher levels in the liver approximately 3-24 hours post-infusion.^8,34^ In our studies, the liver signal drops to baseline at approximately 30-45 minutes following injection, indicating that the cells have moved out of the general circulation and into other tissues around this time. Our approach allows us to visualize early time points post-infusion, providing crucial insights into initial CAR-T cell interactions with tissues. While CAR-T cell efficacy is often assessed at later time points, understanding these early dynamics remains essential for optimizing treatment outcomes, ensuring effective targeting from the outset, and addressing potential challenges such as poor infiltration or early off-target effects. Currently, CAR-T cells are known to accumulate in tissues such as the lung within the first day of therapy, however earlier time points have not been rigorously studied.^7,8,34,35^ Future studies will extend these time points to provide a more comprehensive understanding of CAR-T cell dynamics over longer durations.

**Fig. 4.**
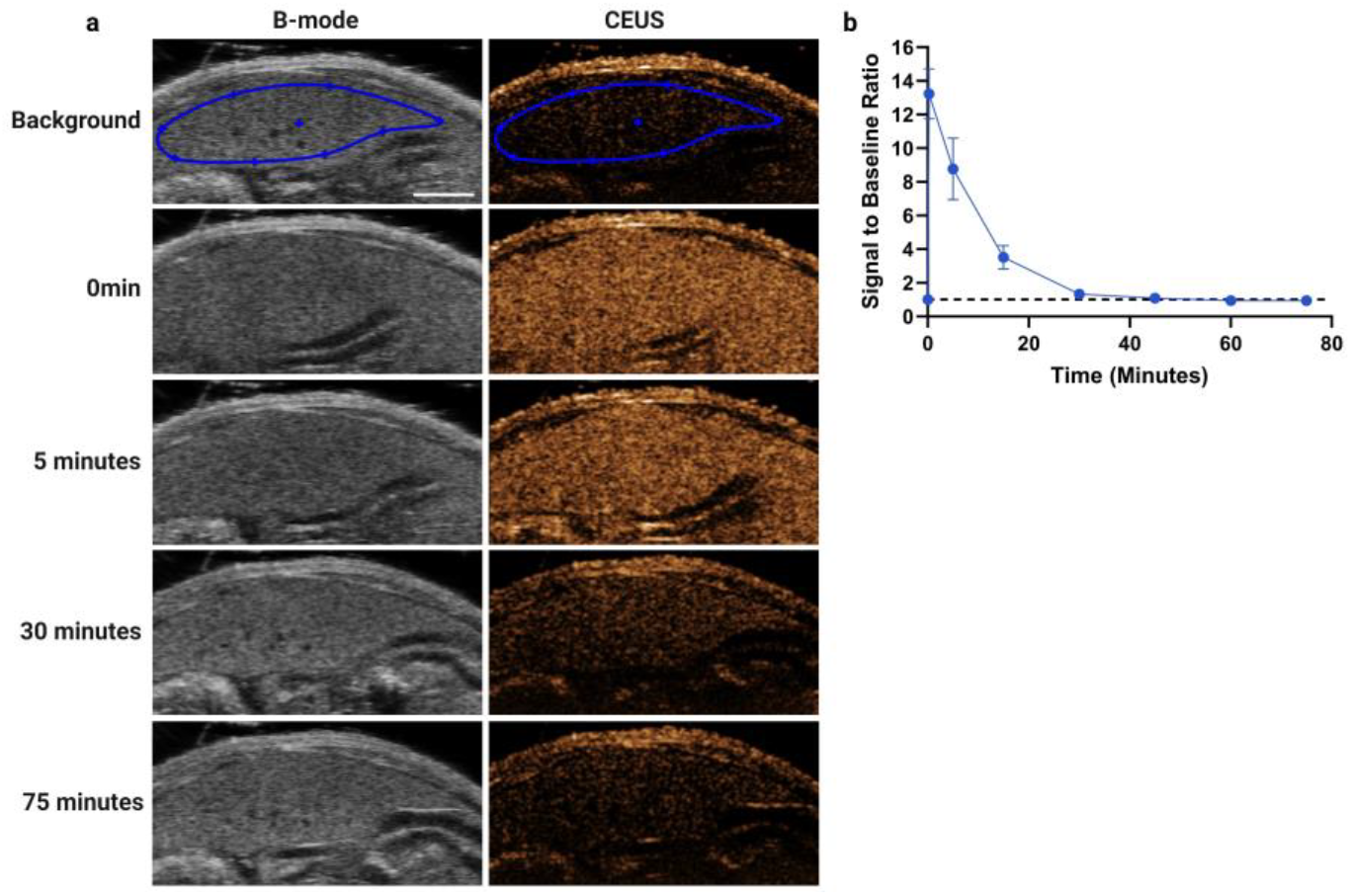
(a) B-mode and CEUS images of the liver of an NCG mouse over 75 minutes before and after the infusion of labeled CAR-T cells (scale bar = 3.5mm). (b) Average signal intensity of the liver over time following the infusion of labeled CAR-T cells (n=3), analyzed using the ROI depicted in the background image of Fig. 4a.

### In vivo detection of labeled CAR-T cells in tumor mouse models

To demonstrate initial proof of concept in a tumor model, NB-labeled CAR-T cells were evaluated in an immunodeficient mouse model using systemic RAJI lymphoma tumors and CD19 CAR-T cells. This model system is beneficial as CD19 CAR-T cells are known to be efficacious against these tumor cells and the CD19 CAR-T cells can traffic to the tumors.^36,37^ In mice, systemic administration of RAJI cells resulted in tumor formation in various organs, but tumors consistently formed in and adjacent to the kidneys. Therefore, we focused our imaging in this region. US imaging was carried out during infusion and at 5, 15, 30, 45, 60, and 75 minutes post-infusion.

Figure 5 shows representative B-mode and CEUS images and analysis for a single mouse. Both tumors adjacent to the kidney and the kidney itself were imaged at each time point. All regions exhibited elevated signal during infusion in the nonlinear mode (Fig. 5b). The variability in image contrast among these regions highlights tissue heterogeneity, which can influence the rate of CAR-T cell infiltration into tumors. This variability is evident in the tumor infusion plot, where regions show different increases in NLC signal. These findings align with published studies highlighting the heterogeneous nature of RAJI tumors.^38-40^ During the initial 30 second infusion, the kidney exhibited a 3-fold increase in contrast suggesting we are detecting the CAR-T cells in circulating transiting through the kidney. Conversely, tumors located near the kidney cortex or adjacent to the kidney itself (denoted as ‘tumor 1’) show reduced contrast likely due to differences in vascular density.

**Fig. 5.**
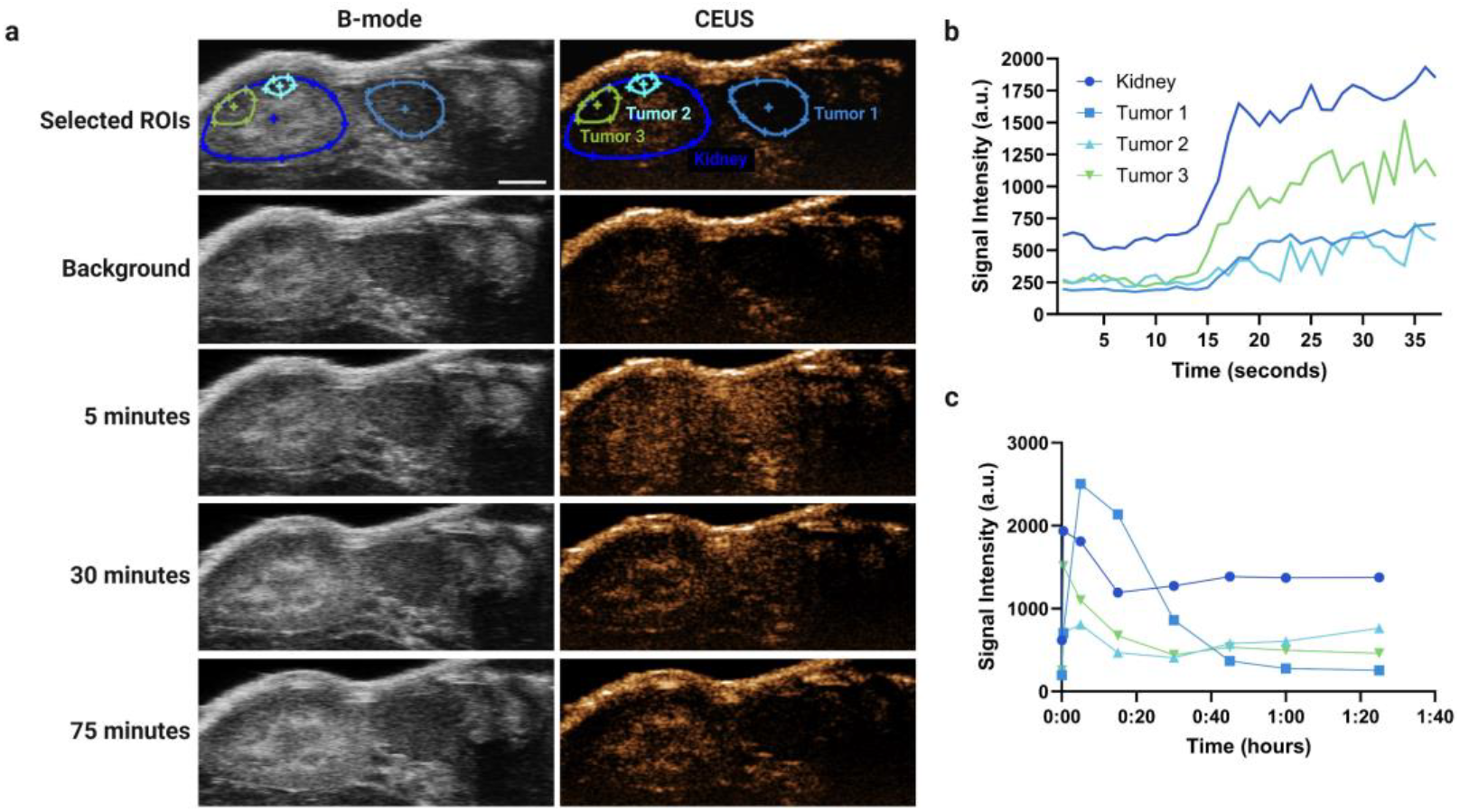
(a) B-mode (L) and CEUS (R) images of the left kidney and tumors of an NCG mouse over 75 minutes. The ROIs (in blue) shown in the images highlight selected areas for analysis in panels (fig. 5b) and (fig. 5c). (scale bar = 2.5mm) (b) Infusion of labeled CAR-T cells into the ROIs depicted in (fig. 5a) for the kidney and tumors. (c) NLC signal intensity within the ROIs at various time points over 75 minutes, where time point 0 represents pre-infusion, and subsequent points are post-infusion of the labeled CAR-T cells.

Following the infusion, images were acquired intermittently over 75 minutes. After 5 min, the signal intensity increased to 5-fold higher levels compared to baseline (Fig. 5c). This latent, but substantial increase in signal suggests a slower but notable cell penetration into the tumor. CEUS images (Fig. 5a) further illustrate the extensive infiltration into the large adjacent tumor at the 5 minute mark. Over time, a spike in NLC signal is observed across all tumors. As the cells become more homogeneously distributed in the blood following the infusion, the signal decreases but stabilizes without dropping below the baseline, indicating persistent but slow CAR-T cell infiltration. Interestingly, ‘tumor 2’ in the kidney shown in Figure 5, shows an increase in signal intensity starting at the 45-minute mark, potentially signaling the onset of cell migration to this area.

Similarly, after the initial spike from the bolus injection and a temporary decrease as the cells distribute uniformly in the bloodstream (while maintaining a signal twice the baseline), the kidney gradually increases in contrast between 15 and 45 minutes. This contrast remains stable from 45 to 75 minutes, signifying ongoing CAR-T cell filtration through the kidney and indicating their persistent presence and potential interaction within the kidney microenvironment.

The mean NLC signal for labeled CAR-T cell distribution in RAJI tumors across four mice is shown in Figure 6. Figure 6a shows post-infusion data normalized by subtracting the background signal. Following infusion, the kidney signal is twice as high as the adjacent tumor signal, underscoring the difficulty CAR-T cells encounter in penetrating and traversing the tumor vasculature. The adjacent tumor signal reaches its peak only 5 minutes after infusion, reflecting the impact of tumor vasculature heterogeneity on CAR-T cell distribution (Fig. 6a). Additionally, the signal-to-baseline ratio for each tissue is illustrated in Figure 6b, underscoring the relative increase compared to the intrinsic NLC signal. While variability between mice can likely be attributed to differences in tumor vasculature and size, similar trends were observed when comparing data from the individual mouse in Figure 5.

**Fig. 6.**
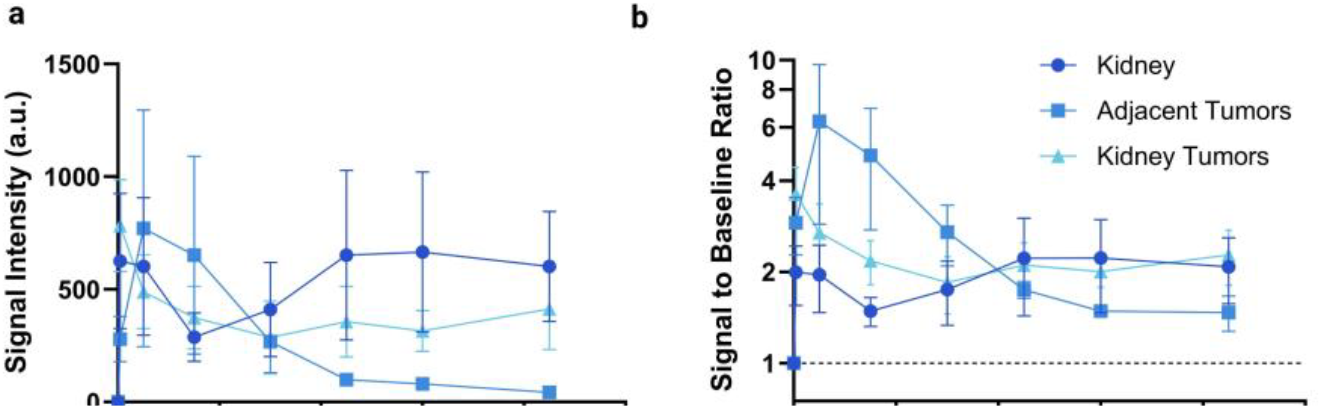
(a) Average NLC signal in tumor bearing mice over 75 minutes. The background was subtracted for each mouse, and the results were averaged (n = 4) (b) The average signal-to-baseline ratio for tumor bearing mice over 75min.

A notable spike in the NLC signal was detected after the bolus injection, which subsequently steadied but remained above baseline. This sustained NLC signal over the 75 minute observation period in the tumor region contrasts with the healthy liver studies, where signals dropped to baseline within 30–45 minutes post-infusion (Fig. 6b). These findings suggest that CAR-T cells can initially infiltrate the tumor region within the first 75 minutes, likely driven by the tumors chemokine signaling, which promotes chemotaxis of immune cells. In comparison, the lack of accumulation in the liver may explain the absence of sustained NLC signal in healthy liver tissue. The liver’s lack of a target antigen and chemokines necessary for CAR-T cell localization is consistent with reports that CAR-T cells do not typically accumulate in the liver until 3-24 hours post-infusion.^8,34-35^ The stabilized NLC signal observed in the kidney tumor region, unlike the transient signal in the liver, indicates CAR-T cell accumulation within the tumor. Given that CAR-T cells typically do not accumulate in healthy kidneys until 1-2 weeks post-infusion in mouse models, the sustained signal above baseline in this study suggests that the presence of CD19 and associated chemokines in the tumor is driving CAR-T cell retention in this region.^8,34^ This early accumulation in the kidney tumor region suggests that the presence of the tumor accelerates CAR-T cell homing and retention.

### Nanobubble labeled CAR-T cell phenotyping and validation

To confirm that NBs were internalized by CAR-T cells, we used rhodamine-conjugated NBs in addition to detecting nonlinear signal via US. Both flow cytometry and confocal microscopy were employed to verify NB labeling (Fig. 7a-b). The presence of intracellular rhodamine signal supports the internalization of the NBs by the CAR-T cells.

**Fig. 7.**
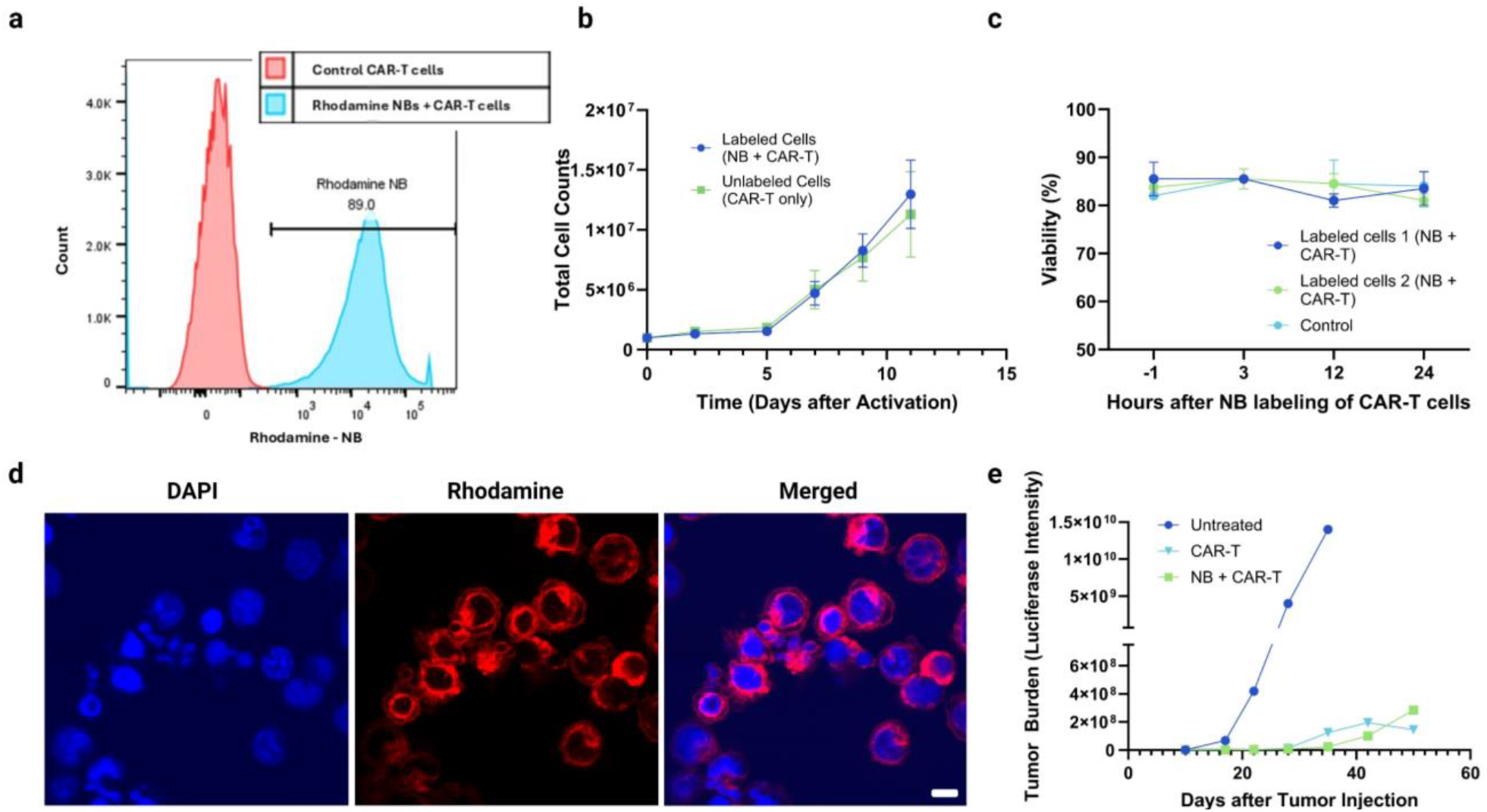
(a) Flow cytometry panel showing the colocalization of CAR expression and rhodamine NB contrast. (b) Proliferation assay comparison of labeled and control CAR-T cells. Expansion showed a non-significant effect on the growth of the cells after activation. (c) Nanobubbles did not impact the viability of the T cells as measured by trypan blue. (d) Confocal microscopy of rhodamine NB-labeled CAR-T cells: red = rhodamine, blue = DAPI (scale bar = 5µm) (e) Mice were intravenously injected with RAJI-luciferase to create tumor burden. Within groups of untreated, control CAR-T, and NB CAR-T, the treated mice both displayed similar levels of survivability, compared to the mice without treatment. The level of tumor burden was measured using luciferase imaging.

To confirm that NB internalization did not affect the phenotype or functionality of CAR-T cells, we compared the proliferation, viability, and cytotoxic activity of the NB-labeled CAR-T cells to unlabeled control cells. As expected, the NB-labeled CAR-T cells expanded similarly to the control cells, with both groups exhibiting an approximately 10-fold expansion over 14 days (Fig 7b; p=0.4703, t-test, ns). Additionally, to verify that the NB labeling process does not impact cell viability, the cells were stained with trypan blue at 0, 3, 12, and 24 hours after NB labeling. Figure 7c illustrates that there was no difference in viability between the labeled and unlabeled CAR-T cells (p=0.77, t-test, ns). This approach allowed for a direct comparison of proliferation rates between labeled and unlabeled CAR-T cells, ensuring that the labeling process did not adversely affect cell growth or viability.

Cy5-conjugated NB-labeled CAR-T cells were compared with unlabeled control CAR-T cells to assess differentiation, with specific attention to markers such as CD45RA, CCR7, and CD27. Differentiation markers, particularly CD45RA and CCR7, remained unchanged compared to the control, indicating that the labeling process did not alter CAR-T cell differentiation (supplementary data). Both CD8 and CD4 populations were analyzed via flow cytometry, showing no appreciable differences in mean fluorescence intensity (MFI) shifts for any of the biomarkers. This suggests a similar distribution of Effector memory (Ems; CD45RA- and CCR7-), Central Memory (CMs; CD45RA- and CCR7+), and terminal effector memory cells (TEMs; CCR7- and CD45RA+) among the labeled CAR-T cells (supplementary data). The viability dye 7-AAD was employed to evaluate potential toxicity and the overall viability of NB-labeled cells relative to the controls. These analyses ensured that NB labeling did not adversely affect CAR-T cell differentiation or viability.

Finally, to further assess the functionality of NB-labeled CAR-T cells, we evaluated their activity in a mouse tumor model. NB labeled and unlabeled CD19 CAR-T cells were injected into immunodeficient mice 1 week after RAJI tumor inoculation. Mice that did not receive any CAR-T cells with disseminated RAJI tumors demonstrated rapid tumor progression, as measured by bioluminescence, and began to succumb to the tumor burden after approximately 5 weeks. In contrast, both NB-labeled CAR-T cells and unlabeled CAR-T cells showed similar disease control (Fig 7e). Our results confirmed no phenotypical or functional differences between the labeled and unlabeled CAR-T cells in terms of proliferation, viability, and cytotoxicity.

## Conclusions

Our evaluation of CAR-T cell therapy, enhanced by NB labeling and US imaging, highlights a promising novel methodology for real-time tracking of CAR-T cell distribution and localization both in vitro and in vivo. This technique has demonstrated efficacy in visualizing CAR-T cell trafficking dynamics, providing insights into therapeutic responses and offering a valuable tool for optimizing treatment strategies. The combination of two FDA-approved techniques highlights the clear translational potential for this procedure to transition into clinical application. In vitro studies confirmed the CAR-T cells are labeled with NBs via flow cytometry and confocal imaging, demonstrating significant contrast under US imaging with a minimum detection threshold of 10^3^ cells/mL, providing a robust method for cell tracking. Additionally, in vivo experiments in healthy mice and RAJI tumor models illustrated the feasibility of monitoring CAR-T cell homing and distribution. The initial bolus injection of labeled cells resulted in a distinct increase in contrast followed by gradual signal decay, reflecting cellular redistribution and infiltration into targeted tissues.

These findings underscore the potential of US imaging with NB-labeled CAR-T cells to transform clinical monitoring, offering insights into biodistribution patterns crucial for treatment efficacy assessment. Future experiments will aim to extend the duration of contrast enhancement in labeled cells and monitor them over longer periods. They will also explore alternative tissues and organs to understand the cells’ distribution better. Moving from in vitro to the clinical application also necessitates optimizing the labeling process, as the costly nature of immunotherapy must ensure high labeling efficiency without compromising efficacy. US NBs are a cost-effective methodology that could be used without hindering the immunotherapy. Additionally, the mechanism of labeling must be investigated to ensure it does not affect the functionality of the cells. Further advancements in this technology could improve personalized therapeutic approaches by optimizing infusion strategies and dosage regimens, thereby enhancing patient outcomes in immunotherapy. While challenges such as signal decay over time and tissue heterogeneity remain, the integration of NB-based US imaging represents a significant step forward in the development of more precise and efficient CAR-T cell therapies for hematological malignancies and potentially beyond, paving the way for future research and clinical applications in cancer treatment. The novelty and importance of being able to track cells continuously after injection, should not be underplayed.

## Acknowledgements

This work was supported by the National Institutes of Health (R21-CA262736) and the Hematopoietic Biorepository and Cellular Therapy Shared Resource of the Case Comprehensive Cancer Center (P30CA043703). The authors thank T. Kosmides for reviewing the manuscript. Figures were created using BioRender.com.

## Conflicts of interest

There are no conflicts to declare.

## Data availability

Data can be provided upon request by reaching out to the authors.

## Notes

### Competing Interest Statement

The authors have declared no competing interest.

