## Supplementary for "In vivo tracking of CAR-T cells in tumors via nanobubble-based contrast enhanced ultrasound"

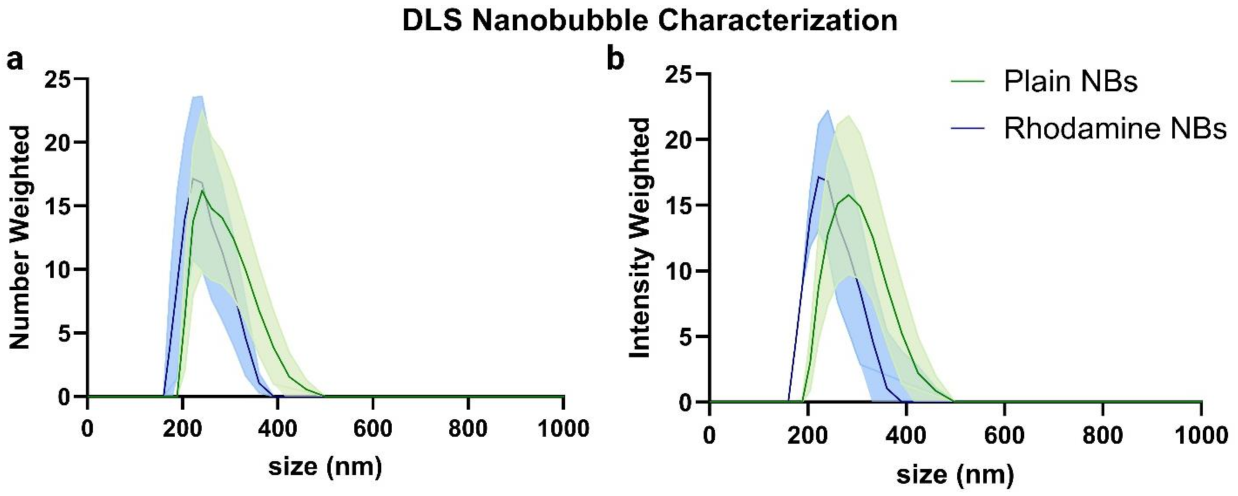


**Fig. S1.** (a) DLS average number-weighted size distribution of plain $C_4F_{10}$ NBs (no fluorophore), n=3. (b) DLS average intensity-weighted size distribution of rhodamine-labeled $C_4F_{10}$ NBs, n=3. The x-axis is limited to 1000 nm because no particles larger than this size were observed.

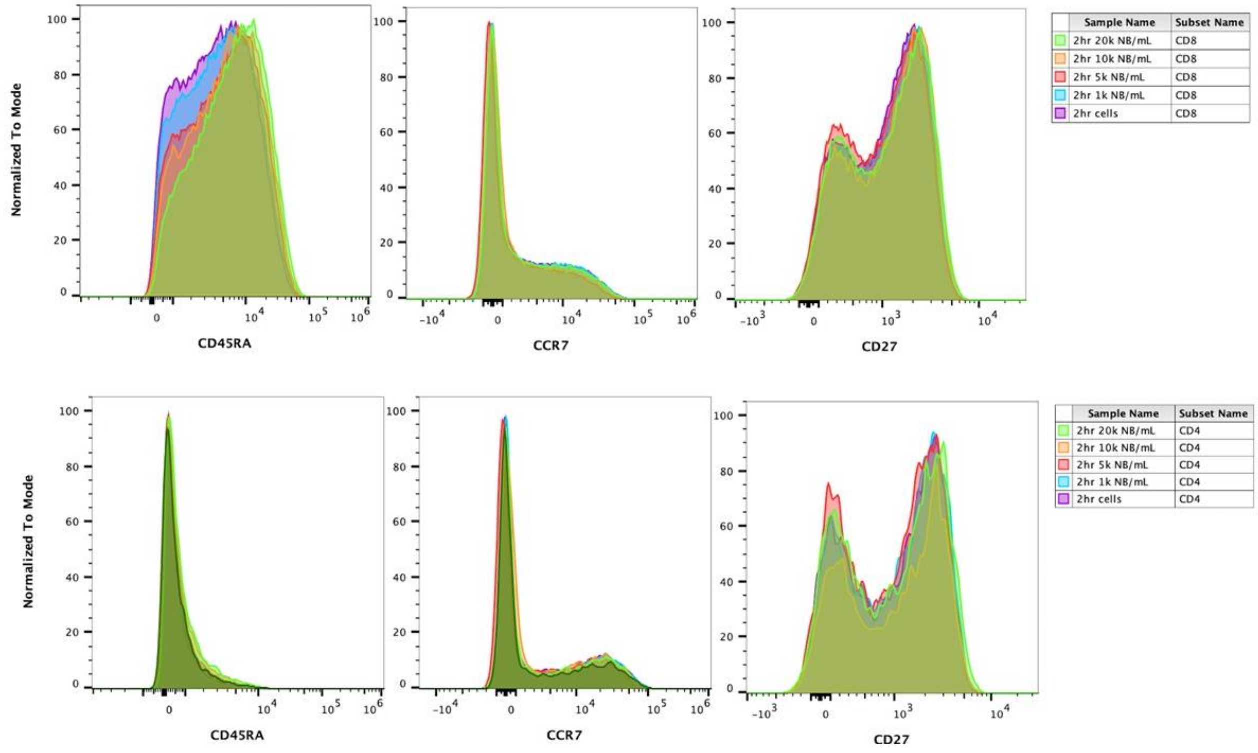


**Fig. S2.** Flow cytometry analysis of CD8+ and CD4+ T cells stained for CD45Ra, CCR7, and CD27 to assess differentiation phenotypes after a 2-hour incubation with varying concentrations of nanobubbles (0, 1k, 5k, 10k, and 20k NBs/mL). No notable shifts in mean fluorescence intensity (MFI) were observed between subsets, suggesting that nanobubble labeling does not impact T cell differentiation across both CD8+ and CD4+ subsets.
